# Spatial point pattern analysis of prehospital naloxone administrations

**DOI:** 10.1101/100412

**Authors:** Christopher Ryan

## Abstract

**Objectives:** The increasing problem in the United States with opioid dependence and overdose, often fatal, is well-recognized. As naloxone has only one clinical use—the treatment of opioid overdose—its administration by EMS personnel can serve as a surveillance indicator for opioid overdose. This study uses specific locations of EMS calls, and methods of point pattern analysis, to detect overall spatial clustering among EMS naloxone administrations compared to EMS calls in general.

**Study Design:** A cross-sectional study of incident locations of EMS responses in a three-county EMS region in the United States.

**Methods:** Repeated random samples from the spatial distribution of all EMS calls were used, in a Monte Carlo simulation, to represent the background inhomogeneity of the population. Observed F, G, and inhomogeneous K and L functions from the spatial distribution of naloxone-involved calls were compared to their null sampling distributions obtained from the Monte Carlo simulation.

**Results:** Cases of naloxone administration demonstrated spatial clustering in the range of 0 to 5000 meters, and particularly around 2500 meters, beyond what could be attributable to the spatial heterogeneity of all EMS calls.

**Conclusions:** Efforts to understand the fundamental nature of opioid overdose as a spatial point process could yield innovative public health interventions to control the epidemic.

## Introduction

The increasing problem in the United States with opioid dependence and overdose, often fatal, is well-recognized. From 1999 to 2008, the age-adjusted annual death rate from prescription opioid analgesics tripled, from 1.5 to 4.5 cases per 100,000 (Centers for Disease Control and Prevention 2011). In a study of state-based mortality files in 28 states, from 2010 to 2012, the age-adjusted annual death rate from heroin overdose doubled, from 1 to 2.1 cases per 100,000 population (Rudd et al. 2014).

A variety of indicators can be used in ongoing surveillance of the opioid dependence problem, and of the corollary problem of opioid overdose. These include death certificates, emergency department visits, hospital admissions, laboratory reports of urine drug tests, and others. Each indicator has its advantages and limitations. Death certificates can be inaccurate, are slow to be processed into final form by centralized vital statistics bureaus, and only capture fatal overdoses. Recovery from opioid overdose can be rapid and complete with proper treatment, rarely requiring hospital admission, rendering admission databases less useful. Since naloxone is a unique and specific opioid antagonist, administration of naloxone by emergency medical services (EMS) personnel is a potentially useful epidemiological indicator of opioid overdose.

Temporal trends in naloxone use can of course be monitored, but spatial patterns may also be of interest. Published studies of the spatial aspects of pre-hospital naloxone administration have tended to use areal or lattice methods, aggregating counts or rates of events by administrative districts such as states, towns, or census tracts. Merchant and colleagues described the temporal and spatial distribution of EMS naloxone administration in the state of Rhode Island between 1997 and 2002, encompassing about 732,000 ambulance runs. They found that per-census-tract rate of naloxone use varied from zero to 3.5% of runs. A few high-incidence census tracts were identified near Providence, the state’s capital and largest city (Merchant et al. 2006). Klimas and colleagues in Dublin, Ireland, took a similar approach and found similar variation in the rate of naloxone-involved EMS calls by administrative area. High-incidence areas were concentrated in the city center (Klimas et al. 2014).

An areal approach is informative but is subject to the well-known modifiable areal unit problem (observed patterns depend upon the size of the administrative unit—census tract, town, county, state—to which the data are aggregated) and to issues of ecological inference (observed patterns among the areal units don’t necessarily apply to the individuals within them). Given that the incident location is usually available in EMS databases, it seems efficient and informative to apply methods for the analysis of spatial point patterns. These methods might yield insights unavailable from an areal approach. I conducted an exploratory analysis of EMS calls for presumed opioid overdose (as indicated by the administration of naloxone) as a spatial point process. Any analysis of the spatial distribution of a human health event must account for the underlying spatial distribution of the susceptible population, which undoubtedly is heterogenous. Furthermore, the spatial distribution of the types of people who tend to call ambulances, and of the locations to which ambulances are called, may differ from that of the population or community as a whole. Thus the main scientific question in this study is whether opioid overdose calls are spatially clustered even after accounting for the heterogeneity of EMS calls in general. In other words, can the observed spatial pattern of opioid overdose calls plausibly be considered a realization of a spatially heterogeneous point process represented by the spatial density of all EMS calls?

## Methods

The setting for this study is an EMS region comprising three counties in southern New York State, with a population of about 300,000 and a surface area of about 6000 *km*^2^. The region comprises rural and suburban areas, a number of villages, and two small cities. It is served by 77 EMS agencies, of various structures: commercial, volunteer, fire-service-based, transporting, non-transporting, advanced life support (ALS), and basic life support (BLS). Intravenous naloxone has been a standard part of ALS protocols in the region for decades. Recently intranasal naloxone has been widely deployed in the region with BLS personnel, firefighters, and police officers as well, but the study period pre-dates that development.

EMS agencies with a total call volume during the study period of less than 50 were excluded, leaving 63 agencies that were invited to participate by allowing their data to be used. Thirteen agencies agreed. These thirteen comprise the largest and busiest agencies, and together they account for over 80% of total regional call volume.

The Regional Emergency Medical Services Council (REMSCO) maintains an electronic database of patient care reports (ePCR) from all EMS agencies in the Region. The database complies with the National Emergency Medical Services Information System (NEMSIS) standards (http://www.nemsis.org/.) Information about incidents and patients is entered into the database by EMS crews almost in real time. The Regional Emergency Medical Advisory Committee (REMAC—a physician subcommittee of the REMSCO) is charged with stewardship of the database and approved its use for this study. The SUNY Upstate Medical University Institutional Review Board also approved the study.

The REMAC provided a data file consisting of incident locations for all EMS calls handled by participating agencies during the study period of 9 September 2012 to 9 February 2014, inclusive. EMS calls in which naloxone was administered to the patient were considered cases, and other calls were considered controls. Incident locations were geocoded in ArcGIS 10.3, using US Census Bureau TIGERline street address files for the three counties, and county boundary shapefiles from the Cornell University Geospatial Information Repository (CUGIR) at http://cugir.mannlib.cornell.edu/. Match accuracy was set at 79%. No manual re-matching of unmatched or tied addresses was attempted. All geographic objects were projected in Universal Transverse Mercator (UTM) zone 18 N with North American Datum (NAD) 1983 and distances in meters. Not unexpectedly, there were a large number of duplicate incident locations among both the cases and the controls. Duplicated locations were removed, meaning that any location to which EMS responded more than once during the study period was represented only once in the data used for analysis.

To assess any human health event for spatial clustering, a representation of the spatial distribution of the underlying human population—the locations in which the event *could* occur—is necessary. In this study, the event of interest is an EMS call in which naloxone was used, while the spatial distribution of all EMS calls represents the universe of possible event locations from which Monte Carlo sampling can be performed.

Two methods of Monte Carlo simulation, in broad outline similar to methods of Moller and Waagepetersen (2004), Waller (2009), Diggle (2014), and Illian (2008), suggest themselves, and both were used in the present study. Each must be considered in light of the fact that, operationally, incident locations are recorded in the EMS database as street addresses which, when geocoded, put all of them within 6 meters of a road centerline. Even if a patient is found in the barn on his farm, the incident will be recorded as the address on his mailbox, hundreds of meters or more way.

In the first method, 500 samples of locations, each equal in size to the number of observed naloxone incidents, were drawn from among the observed locations of all incidents. This has the advantage that all resulting simulated locations are realistic locations for an EMS call, since one has in fact already occurred there. In the second method, a kernel density estimate based on the locations of all EMS calls was obtained, with bandwidth automatically selected via a likelihood cross-validation method (Baddeley, Rubak, and Turner 2016, p. 171). The selected bandwidth was adjusted further, to produce a two-dimensional density that, when plotted, was consistent visually with the observed distribution of incident locations. Then 500 repeated samples, each equal in size to the observed number of naloxone cases, were drawn from that spatial density. This method has the advantage that it includes simulated locations where an EMS call *could* conceivably occur but did not happen to occur during the finite study period. A difficulty with the second method is that it can lead to the spurious appearance of clustering (Baddeley, Rubak, and Turner 2016, p. 735). To overcome that problem, each simulated point drawn from the two-dimensional density was adjusted by projecting it to the nearest road segment. The road network in the study region is shown in Figure 1.

**Figure 1:**
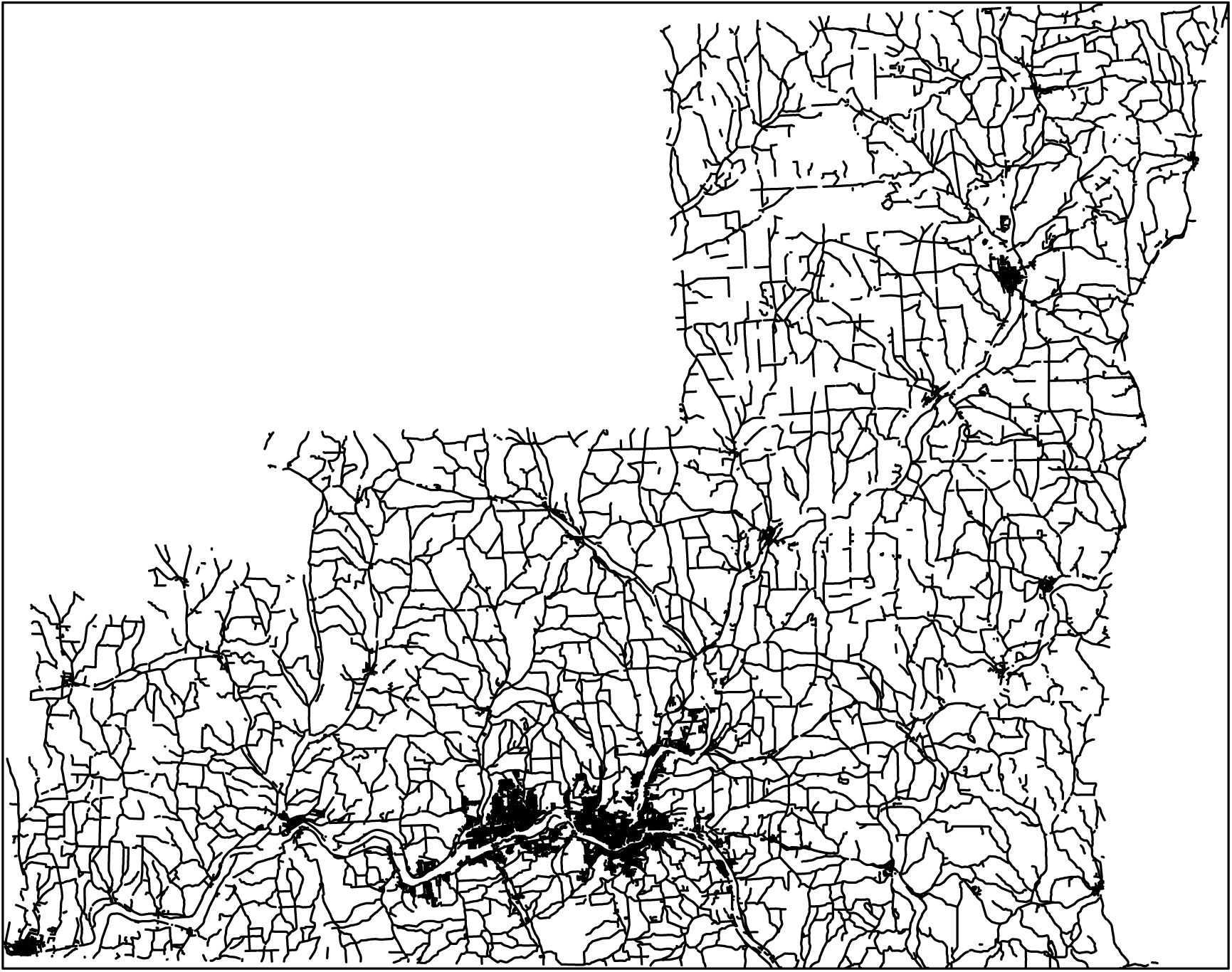
Road network in the Region.

The samples of simulated case locations were used to obtain sampling distributions for the F, G, and inhomogeneous K functions for all EMS calls in general. The inhomogeneous L function was obtained from the inhomogeneous K function via 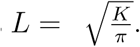 Those same functions derived from the observed case locations alone were then compared graphically to the sampling distributions, testing the null hypothesis that the observed case locations are simply one realization of a spatial point process that generates all EMS calls.

The analysis was conducted in R (R Core Team 2015), relying heavily on the spatstat package (Baddeley and Turner 2005). Where incident locations are displayed, they are jittered within a 500 meter radius and the map scale kept small, to protect privacy; all quantitative analyses, however, were performed using the actual locations.

## Results

After eliminating unmatched addresses and reducing duplicated locations to single points, the analytical dataset consisted of 183 cases and 10,643 controls (Table 1). Their approximate locations are shown in Figure 2, where the expected heterogeneity is obvious.

**Table 1.**
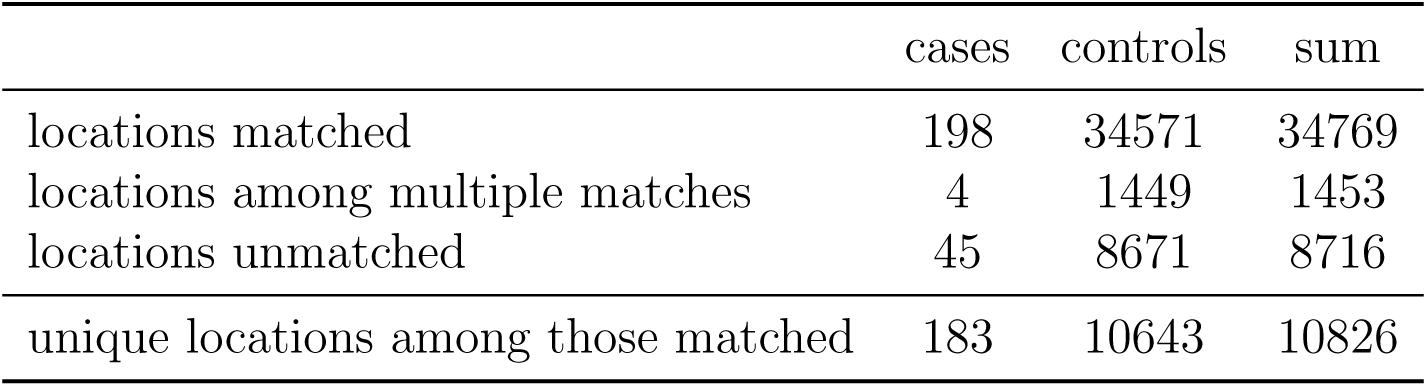
Unmatched, tied, and duplicate locations, and the final analytical dataset.

|  | cases | controls | sum |
| --- | --- | --- | --- |
| locations matched | 198 | 34571 | 34769 |
| locations among multiple matches | 4 | 1449 | 1453 |
| locations unmatched | 45 | 8671 | 8716 |
| unique locations among those matched | 183 | 10643 | 10826 |

**Figure 2:**
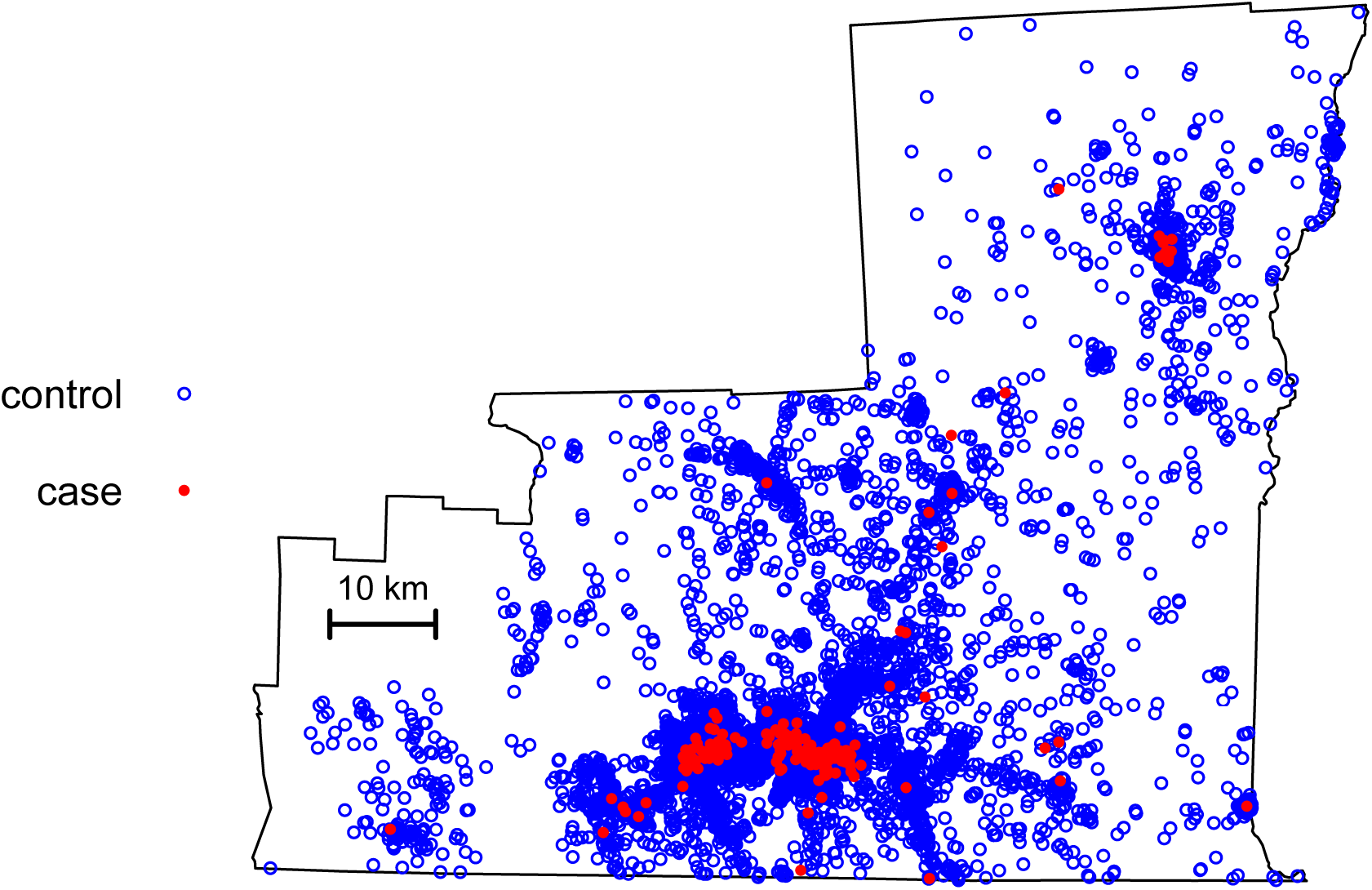
Approximate locations of naloxone cases and non-naloxone controls.

### Monte Carlo sampling from observed incident locations

Five hundred samples, each of size 183 (the observed number of unique naloxone-involved incidents), were drawn from the set of all 10,643 unique and matched incident locations. These five hundred samples yielded sampling distributions for the G, F, and inhomogeneous K and L functions, against which those same functions derived from the observed naloxone incident locations could be compared. For illustrative purposes, four of the five hundred Monte Carlo samples are shown in Figure 3.

**Figure 3:**
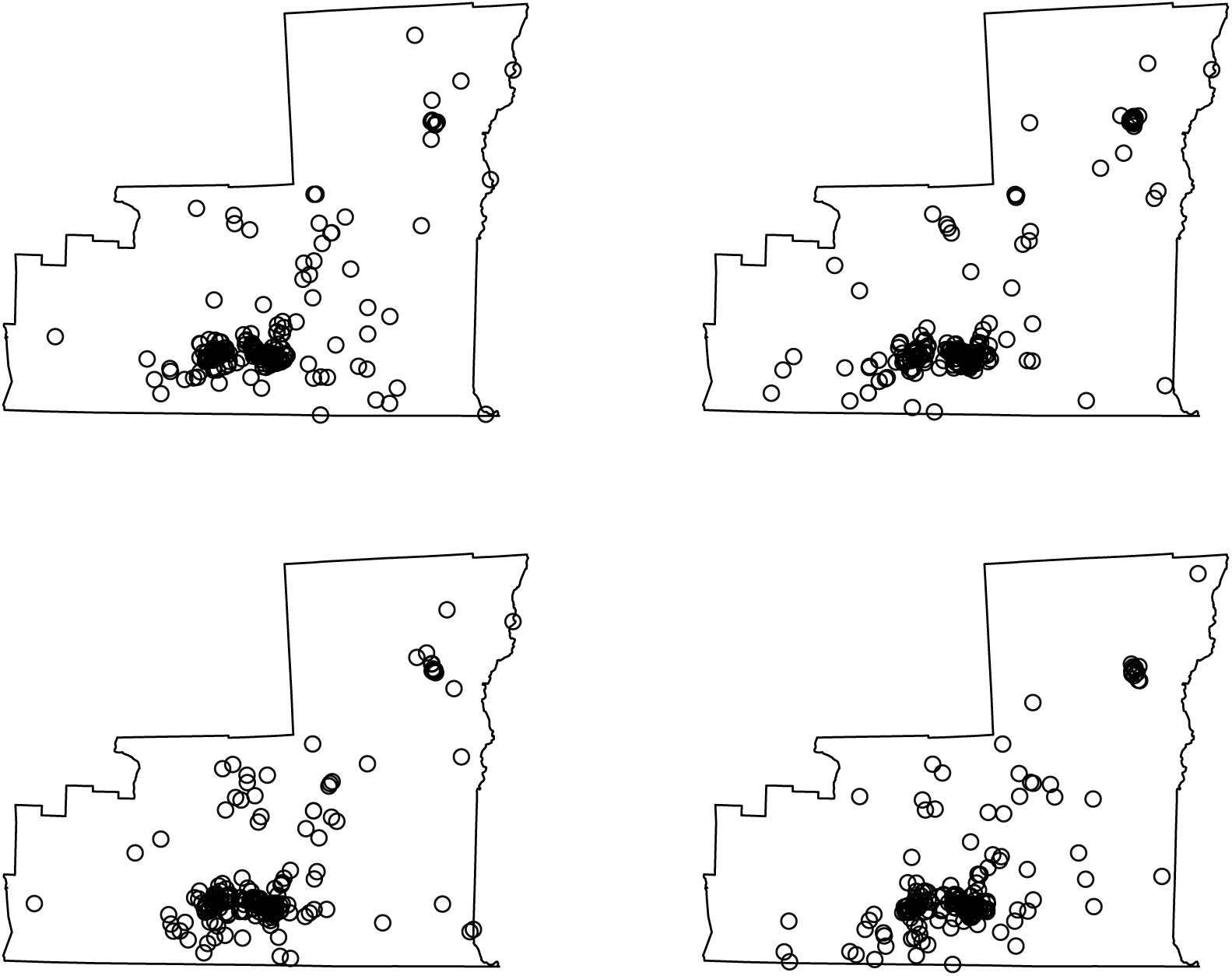
Four of the 500 simulated point patterns produced in the Monte Carlo simulations from actual EMS call locations, for comparison with the observed pattern of all cases and of controls shown in Figure 2.

Four graphical assessments of spatial clustering, the F, G, and inhomogeneous K and L functions are shown in Figure 4. The G and L functions of the observed naloxone cases suggest that shorter inter-event distances are overrepresented among the naloxone calls, compared to inter-event distances among all EMS calls in general. The F function from the observed naloxone cases suggests that smaller event-free spaces are underrepresented among the naloxone calls, compared to the event-free spaces among all EMS calls in general. Taken together, these observations are consistent with spatial clustering, most prominent in a radius of about 2500 meters, of naloxone calls over and above that seen in EMS calls in general

**Figure 4:**
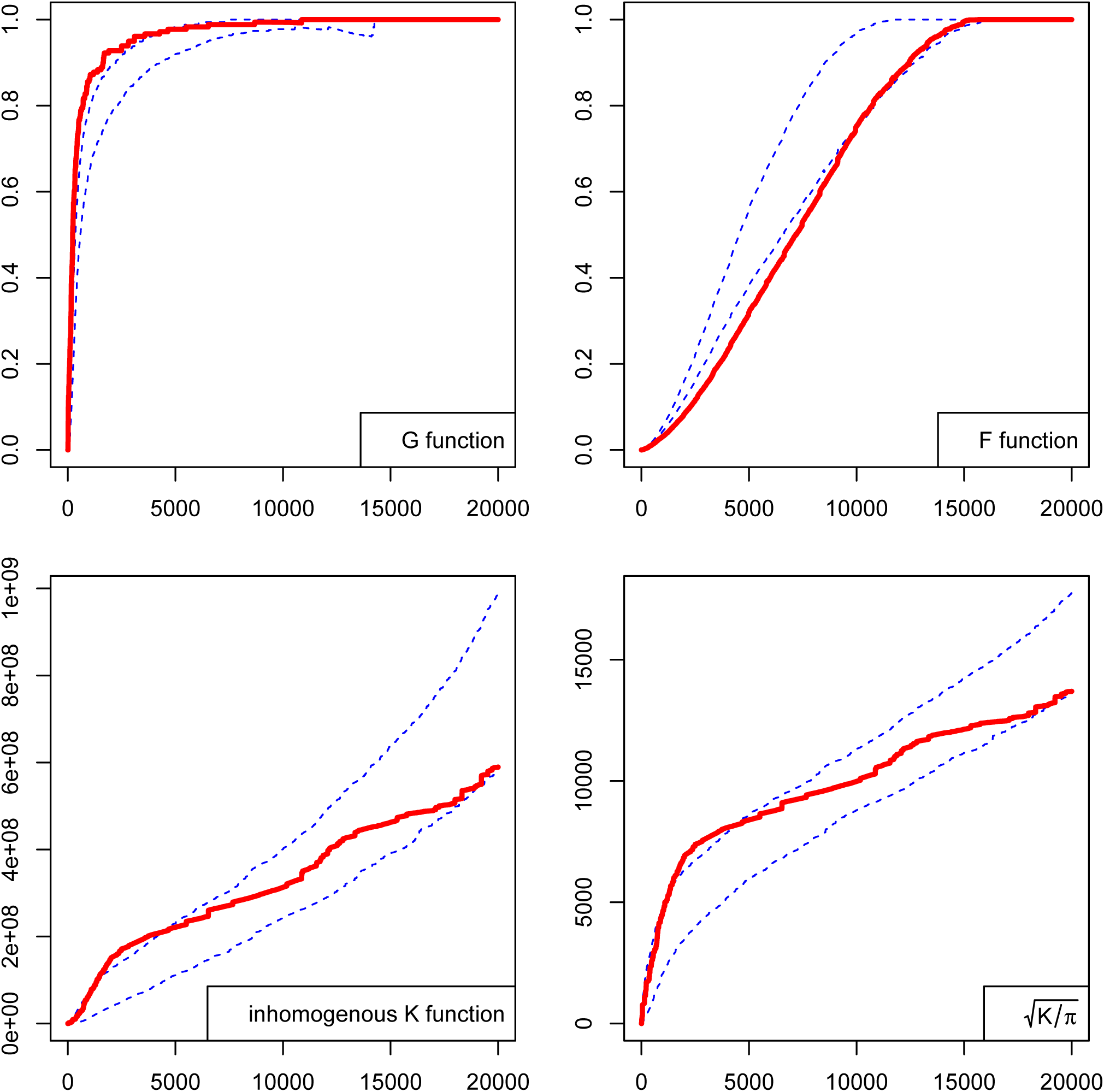
Four measures of spatial clustering of naloxone-involved EMS calls compared to that of EMS calls in general. Heavy lines are the functions from the 183 observed naloxone incidents. Dotted lines are the bounds of pointwise 95% acceptance intervals obtained from Monte Carlo simulation with points drawn from the actual locations of all EMS calls.

#### Monte Carlo sampling from a two-dimensional spatial density

The two-dimensional density derived from the observed locations of all incidents, and covering the entire study region, was used to draw five hundred samples, each of size 183 (the observed number of unique naloxone-involved incidents). Each simulated point in each sample was displaced to the nearest road segment, to mimic the way in which incident locations are recorded in the EMS database. Table 2 describes the distances by which the 91500 simulated case locations (500 simulations of 183 points each) needed to be displaced, to reach the nearest road segment. For illustrative purposes, four of the five hundred Monte Carlo samples are shown in Figure 5.

**Table 2:** Summary of distances, in meters, by which simulated case locations were moved, to reach the nearest road.

|  | distance |
| --- | --- |
| minimum | 0.0 |
| 25th percentile | 31.3 |
| median | 98.6 |
| 75th percentile | 246.1 |
| maximum | 1964.2 |

**Figure 5:**
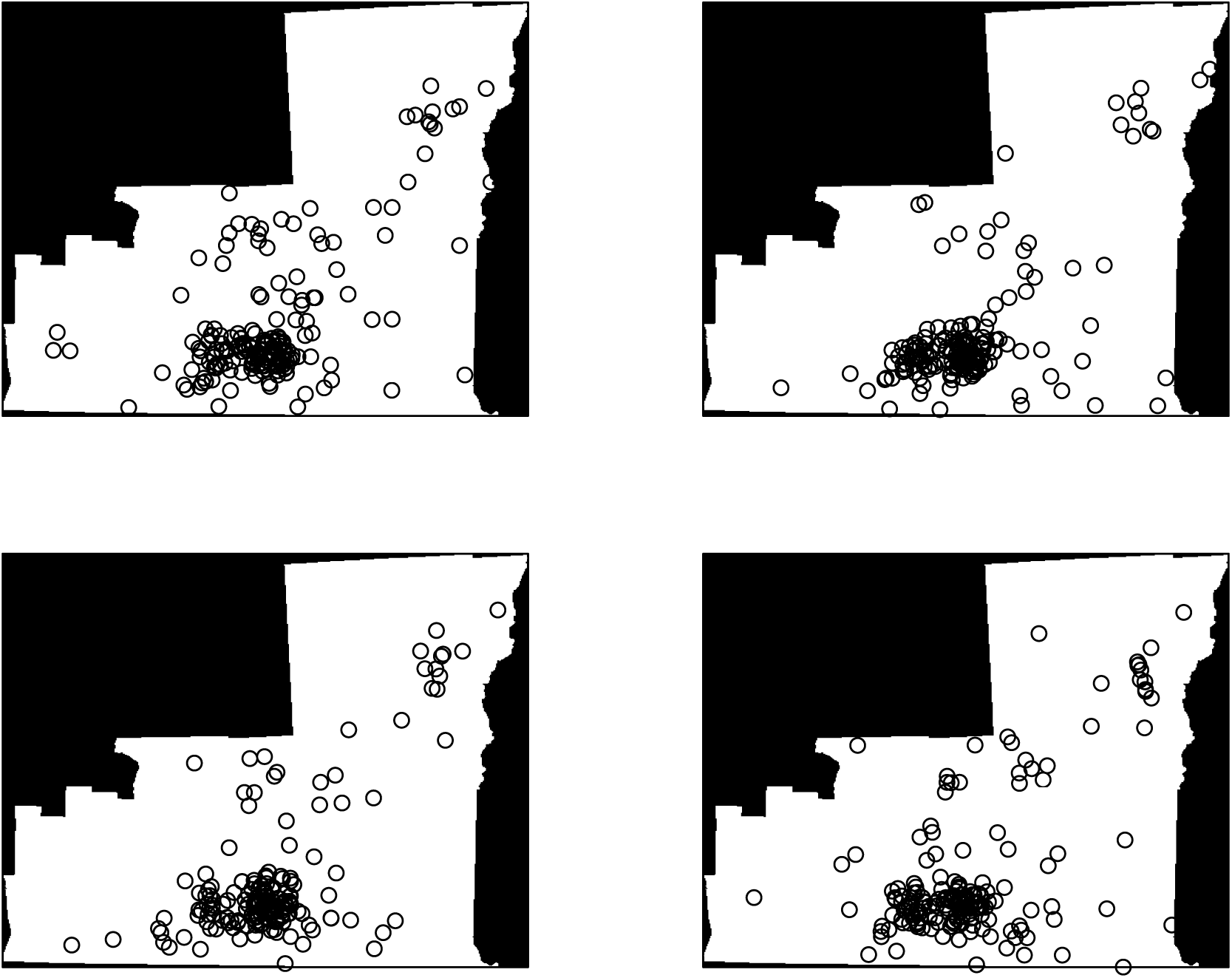
Four of the 500 simulated point patterns produced in the Monte Carlo simulations from a two-dimension density derived from all EMS call locations, for comparison with the observed pattern of all cases and of controls shown in Figure 2.

These five hundred samples yielded sampling distributions for the G, F, and inhomogeneous K and L functions, against which those same functions derived from the observed naloxone incident locations could be compared. The graphical assessments of spatial clustering, shown in Figure 6, again suggest spatial clustering of naloxone-involved EMS calls ompared to all EMS calls in general, especially around a radius of 2500 meters.

**Figure 6:**
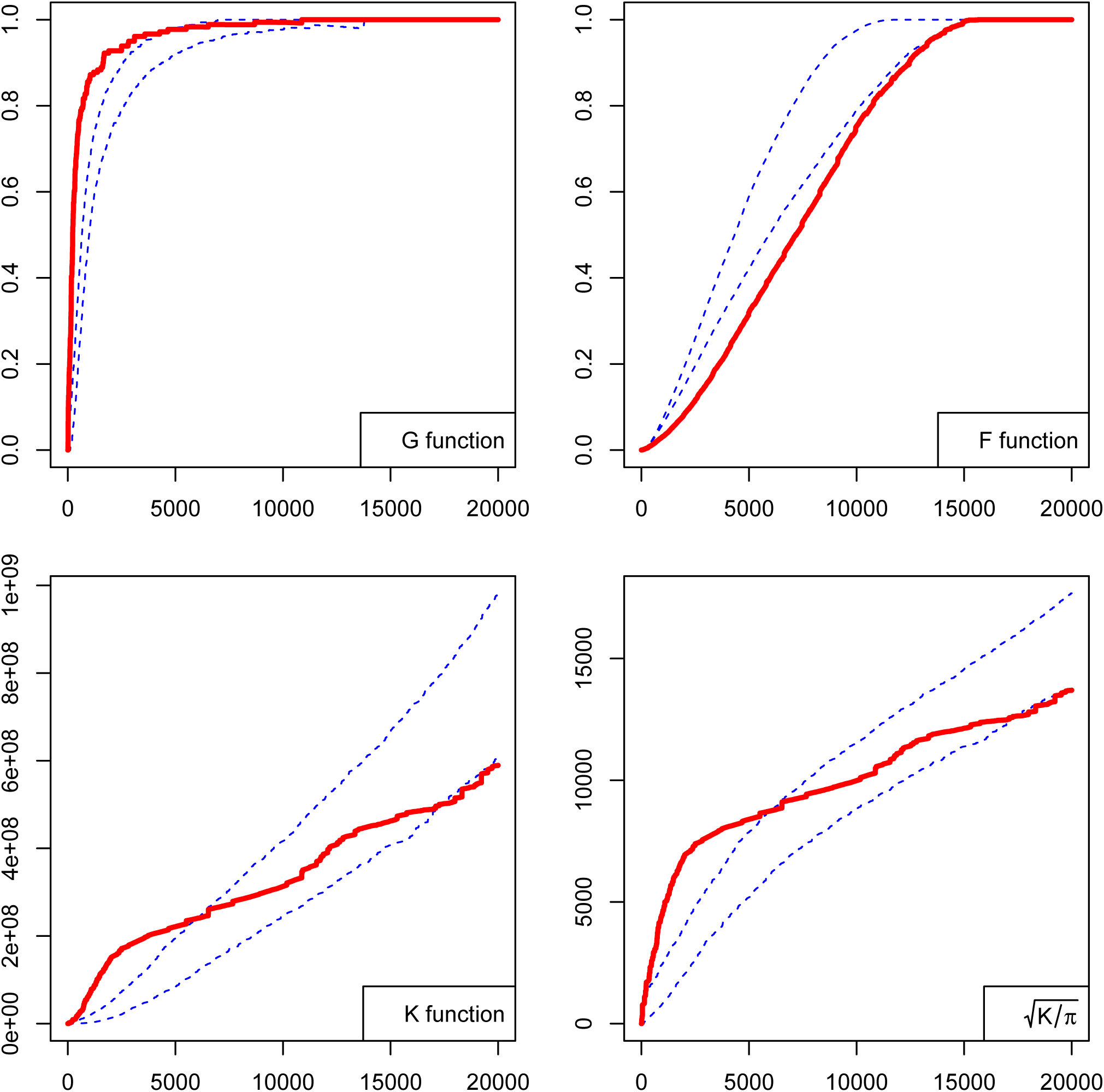
Four measures of spatial clustering of naloxone-involved EMS calls compared to that of EMS calls in general. Heavy lines are the functions from the 183 observed naloxone incidents. Dotted lines are the bounds of pointwise 95% acceptance intervals obtained from Monte Carlo simulation with points drawn from a two-dimensional spatial density representing the distribution of all EMS calls.

## Discussion

The results of this point pattern analysis of the locations of EMS calls are consistent with spatial clustering of presumed opioid overdose calls, beyond that to be expected among EMS calls in general. Clustering was most evident in a range of around 2500 meters. This supports, via the methods of point pattern analysis, similar findings from methods for lattice or areal data (Klimas et al. 2014; Merchant et al. 2006).

This was a study of global clustering of naloxone-involved EMS calls, not a search for specific clusters. The distinction is important and is explained well by Besag and Newell: “In tests of clustering, the usual question is whether the observed pattern of disease in one or more geographical regions could reasonably have arisen by chance alone.” In the present study, the null hypothesis of “chance alone” is adjusted for the spatial distribution of all EMS calls. In contrast, the purpose of cluster detection is to search, perhaps repeatedly, “…a very large expanse of small administrative zones for evidence of individual ‘hot spots’ of disease…” (Besag and Newell 1991). Both strategies are useful, but for different purposes. The global clustering found in this study is perhaps best interpreted as justification for more complex modeling of the pattern-generating process, so as to understand the reasons for spatial clustering of opioid overdoses. For example, first-order effects would include various physical, demographic, or socioeconomic characteristics of neighborhoods that might attract opioid users or overdosers. Second-order effects imply interaction between points—that the occurrence of one opioid overdose in a particular location influences other people to overdose nearby. This descriptive study cannot offer explanations, and disentangling first-from second-order effects is often difficult; both may be at work simultaneously. Understanding the relative contributions of various first- and second-order effects implied in different point process models could guide public health mitigation strategies.

In the analysis of spatial point patterns of human health events, it is often difficult to find an appropriate comparison population—a representation of the spatial distribution of the humans to which that health event pertains. This study capitalized on the availability of a large, comprehensive, three-county ePCR database that provided locations of all EMS calls. This reference population served to control for the expected inhomogeneity in the spatial distribution of the regional population, and for spatial inhomogeneity in the propensity for people from different neighborhoods to call ambulances—perhaps related to socioeconomic status, distance from hospitals, availability of transportation, family support, and other factors.

The representation of locations as street addresses presented a challenge. However, projecting the Monte Carlo sampled locations to the nearest road is a reasonable solution and has the advantage of being consistent with real-world EMS practice.

Several limitations must be considered. Participation by EMS agencies was not universal; however, the vast bulk of EMS calls in the region was included. Duplicate locations were eliminated, mainly to justify the use of relatively simple methods to detect clustering. Thus the analytical data set comprised locations where an ambulance had responded at least once during the study period. Methods that accomodate duplicate locations may yield different results. Automated geocoding was incomplete, with many locations left unmatched. While manual matching could have been undertaken with the unmatched case locations, it would have been impractical to do the same for the much larger number of unmatched control locations (see Table 1). The methodologic rigor of treating cases and controls in the same manner outweighed any value to be obtained from manually matching the remaining case locations to enable their inclusion in the dataset. The Figures show point-wise, not global, acceptance intervals and thus must be interpreted with some caution. Lastly, naloxone administration is a useful but imperfect indicator of opioid overdose. Some patients suffering opioid overdose do not receive naloxone—for example, patients obviously dead when EMS arrives. Conversely, a small number of patients not suffering from opioid overdose will in fact be given naloxone, based on the paramedic’s best judgment at the scene.

From a broader perspective, this study presents an example of using a regional EMS database to study spatial aspects of a public health problem. With increased use of NEMSIS-compliant databases, and with incident location almost universally recorded as a routine part of EMS operations in the field, these data can become increasingly useful in studying spatially-varying health events.

## Acknowledgements

I thank the leadership and field providers of the Susquehanna Emergency Medical Services Region for their support.

